# Bacterial invasion across the human skin barrier – mechanisms and ensuing tissue degradation

**DOI:** 10.1101/2021.10.06.463452

**Authors:** Zachary W. Lipsky, Marisa Patsy, Cláudia N. H. Marques, Guy K. German

**Affiliations:** Department of Biomedical Engineering, Binghamton University, 4400 Vestal Parkway East, Binghamton, NY 13902.; Department of Biological Sciences, Binghamton University, 4400 Vestal Parkway East, Binghamton, NY 13902.; Binghamton Biofilm Research Center, Binghamton University, 4400 Vestal Parkway East, Binghamton, NY 13902.; Fischell Department of Bioengineering, University of Maryland, 3102 A. James Clark Hall, College Park, MD 20742

**Keywords:** Stratum Corneum, *Staphylococcus aureus*, Lipids, Mechanics

## Abstract

Atopic Dermatitis (AD) is associated with a deficiency of skin lipids, increased populations of *Staphylococcus aureus* in the microbiome, and structural defects in the stratum corneum (SC), the outermost layer of human skin. However, the pathogenesis of AD is ambiguous as it is unclear whether observed changes are the result of AD or contribute to the pathogenesis of the disease. Previous studies have shown that *S. aureus* is capable of permeating across isolated human SC tissue when lipids are depleted to levels consistent with AD conditions. In this study, we expand upon this discovery to determine the mechanisms of bacterial penetration into the SC barrier. Specifically, we establish whether bacteria are permeating intercellularly, between corneocytes, or employing a combination pathway of both inter- and intra-cellular travel. The mechanical implications of bacterial invasion, lipid depletion, and media immersion are also evaluated using a newly developed, physiologically relevant, temperature-controlled drip chamber. Results reveal that *S. aureus* can be internalized by corneocytes, indicating transcellular movement through the tissue during permeation, consistent with previous theoretical models. *S. aureus* also degrades the mechanical integrity of human SC, particularly when the tissue is partially depleted of lipids. These observed mechanical changes are likely the cause of broken or ruptured tissue seen as exudative lesions in AD flares. This work further highlights the necessity of lipids in skin microbial barrier function.

**Importance:** Millions of people suffer from the chronic inflammatory skin disease Atopic Dermatitis (AD), whose symptoms are associated with a deficiency of skin lipids that exhibit antimicrobial functions, and increased populations of the opportunistic pathogen *Staphylococcus aureus*. However, the pathogenesis of AD is ambiguous, and it remains unclear if these observed changes are merely the result of AD, or contribute to the pathogenesis of the disease. In this article, we demonstrate the necessity of skin lipids in preventing *S. aureus* from penetrating the outermost barrier of human skin thereby causing a degradation in tissue integrity. In terms of AD, this bacterial permeation into the viable epidermis could act as an inflammatory trigger of the disease and could also explain tissue fragility and lesion formation seen with AD patients. Moreover, bacterial induced degradation could lead to increased pathways and further allergen intervention creating chronic irritation.

## INTRODUCTION

Millions of children and adults suffer from atopic dermatitis (AD) (1, 2), a chronic inflammatory skin disease. AD symptoms are associated with a deficiency of skin lipids (3), increased populations of *Staphylococcus aureus* in the microbiome (4), and structural barrier defects in the stratum corneum (SC) (5), the most superficial layer of human skin. However, the pathogenesis of AD is ambiguous, and it is unclear if these observed changes are the result of AD or contribute to the pathology of the disease (6). We hypothesize that decreases in SC lipid populations may cause sufficient barrier dysfunction to enable *S. aureus* to permeate across the epidermis, and act as a potential inflammatory trigger of the disease. This could explain the subset of cases that do not fit a genetically motivated disease development, either with mutations in immune associated genes (IL4, IL13, RANTES, CD14, NOD1) (7, 8) or barrier function associated genes (FLG) (7, 9, 10). Our previous work has shown that *S. aureus* is capable of permeating across isolated human SC tissue when lipid populations decrease to levels consistent with AD conditions (11). As an extension of this work, we aim to utilize a newly developed physiologically relevant temperature- and moisture- controlled drip chamber to investigate the pathways that allow *S. aureus* to permeate across the tissue and the mechanical and structural implications of bacterial invasion.

## RESULTS

### Lipid depletion allows permeation of *S. aureus* through the SC and internalization into corneocytes

The ability of *S. aureus* to penetrate isolated healthy unaltered control and lipid depleted (Delipid) human SC is first assessed. Three-dimensional bacterial penetration depths within SC samples were characterized over a 5-day period. Fig. 1 shows representative composite fluorescent profile cross sections through the SC, with *S. aureus* shown in green and free fatty acids within the SC shown in red. Figs. 1A and B show the position of bacteria on day 0, 2 hours after inoculation for control and lipid depleted SC samples respectively. Figs. 1C and D show bacterial positions on day 4 for the same respective samples. Fig. 1D shows that bacteria can permeate into the lipid depleted SC, while Fig. 1C shows that bacteria on control SC do not, consistent with prior studies(11).

**Figure 1.**
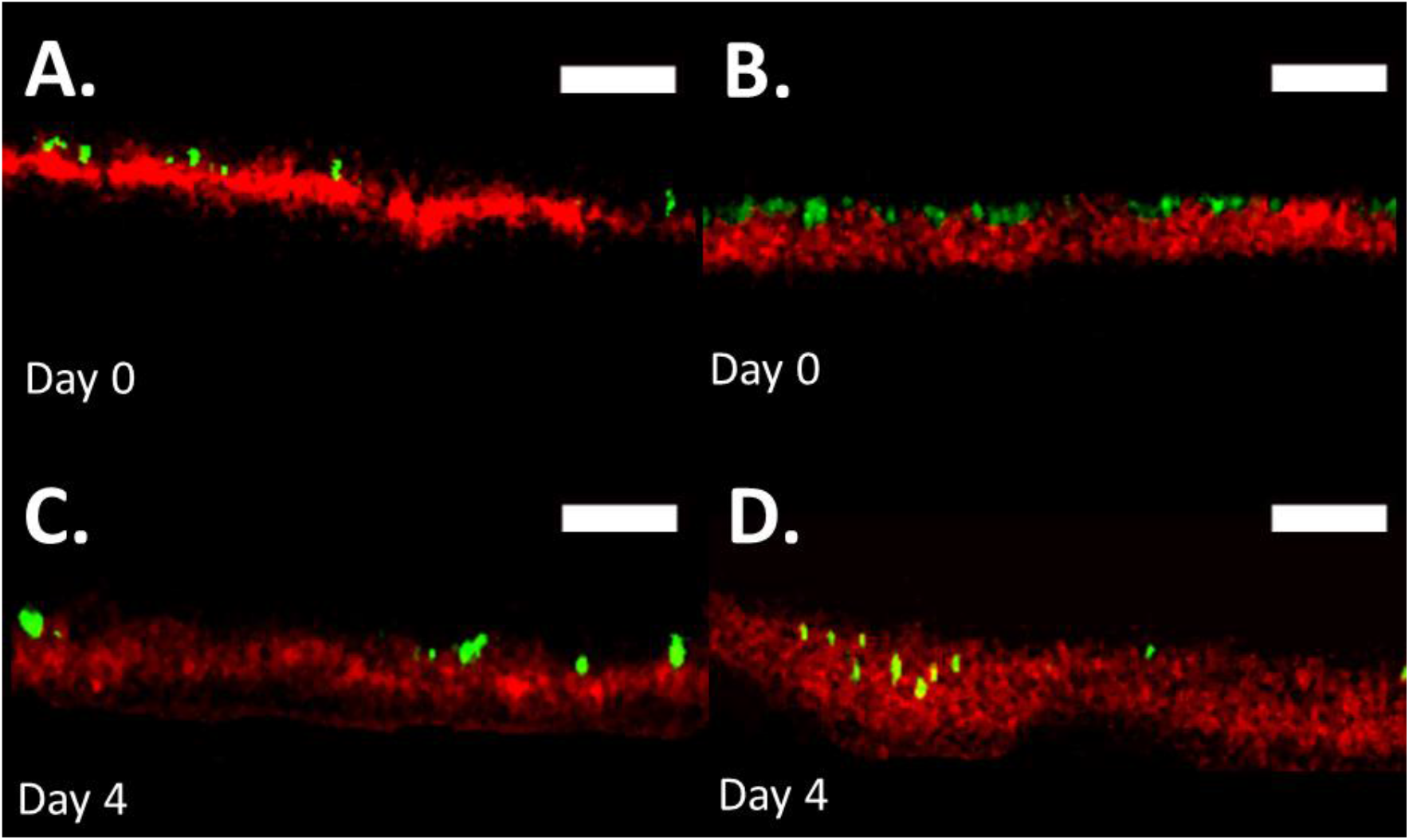
*S. aureus* penetrates lipid depleted SC in drip chamber setup. Fluorescent confocal cross-sectional profiles of *S. aureus* permeation into a control (A & C) and delipidated (B & D) SC sample over a 5-day period. Images show representative fluorescent cross-sectional profiles of BODIPY lipid stained SC (*red*) inoculated with GFP labeled *S. aureus* bacteria (*green*) after (A & B) 0, and (C & D) 4 days. An identical contrast change has been imposed on all images in each lipid condition to enhance visual clarity. *Scale bar* – 15 μm.

The interactions between bacteria and the delipidated SC tissue microstructures are further investigated to better understand the permeation pathways. Previous reported models of permeation highlight that purely intercellular permeation pathways are unlikely compared with combined inter- and intra-cellular permeation, due to permeation timescales being notably larger than those observed (11). Permeation of these non-motile bacteria is modeled as a 3D random walk process, where the position of a new bacteria formed through binary fission is located arbitrarily next to the parent cell. For a combined inter- and intra-cellular permeation pathway to be viable however, *S. aureus* must utilize a direct transcellular route through corneocytes. Internalization of *S. aureus* has previously been shown to occur both *in vivo* and *in vitro* within keratinocytes at various levels of differentiation up until the *stratum granulosum* (12, 13). However, internalization in anucleated corneocytes that exhibit a cornified envelope (14, 15) has not previously been observed. As such, bacterial internalization assays are next performed using methods described in Kintarak *et al.*(16).

Fig. 2A shows the averaged (11 ≤ *n* ≤ 15 independent samples for each condition) *S. aureus* population in CFU/mL on delipidated and control SC samples partially embedded in an ethylene-vinyl acetate substrate (EVA), and EVA substrates alone. For each condition populations are quantified with and without a gentamicin treatment after a 5-day growth period. Population data is normalized and scaled to the greatest CFU/mL magnitude per trial (*n* = 4). There are no significant differences between conditions without the addition of gentamicin, except between EVA alone and control SC. As such, bacterial populations here are not statistically affected by SC lipid concentration over the 5-day timescale. As expected, bacterial population viability reduces when treated with gentamicin. Fig. 2B shows a rescaled bar chart from Fig. 2A. For gentamicin treated conditions, the reduction of bacterial population is significantly smaller in delipidated SC tissue samples, while no statistical difference exists between gentamicin treated EVA substrates and control SC samples.

**Figure 2.**
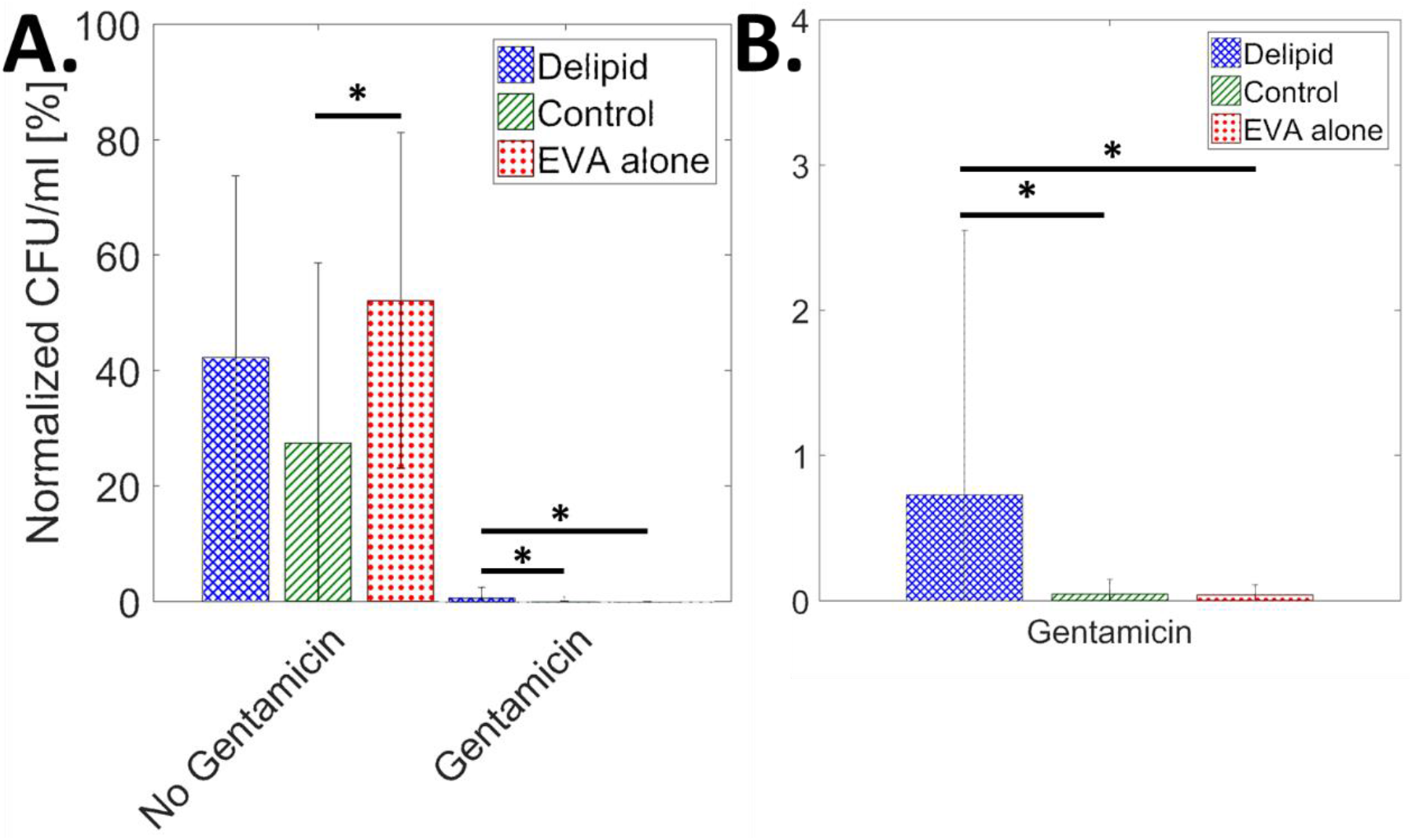
*S. aureus* is internalized by corneocytes in lipid depleted SC. (A) Average normalized bacterial population in CFU/mL, from *n* = 4 independent trials, scaled by the highest CFU/mL per trial, for delipidated SC (*blue crosshatch*), control SC (*green stripes*), and EVA alone (*red dots*), with and without the addition of gentamicin (11 < *n* < 15 for each condition). (B) Averaged normalized CFU/mL rescaled for gentamicin effected conditions only.

### *S. aureus* alters SC mechanical properties

We further examine the effect that *S. aureus* colonization, growth, permeation, and cellular internalization has on the mechanical properties of control and delipidated SC. To assess the independent effects of media immersion and bacterial colonization and permeation, SC samples are exposed to either *S. aureus* and media combined, or media alone. These are then compared with unexposed controls. For each condition, all SC samples are either equilibrated for 24 hr to 25 or 100% relative humidity (RH) prior to uniaxial testing to examine the impact of the exposure to the linear elastic and plastic regimes of the tissue. The isolated effect of lipid depletion is further established by comparing control and delipidated SC mechanical properties without exposure to media, while the isolated effect of media immersion is determined for each lipid condition by comparing SC exposed to media with non-immersed counterparts.

Fig. 3 shows the average elastic modulus, *E*, fracture stress, *σ_f_*, fracture strain, *γ_f_*, and work of fracture, *W_f_* of control and delipidated SC subjected to the various treatment conditions, then equilibrated to 25% RH. Fig. 3A shows that the elastic modulus increases only resulting from lipid depletion, consistent with previous studies(17). Neither media immersion nor bacterial growth alter the stiffness of the tissue. Figs. 3B–D further highlight that at low humidity conditions, neither lipid depletion, media immersion, nor bacteria have a significant effect on the fracture stress, fracture strain, or work of fracture.

**Figure 3.**
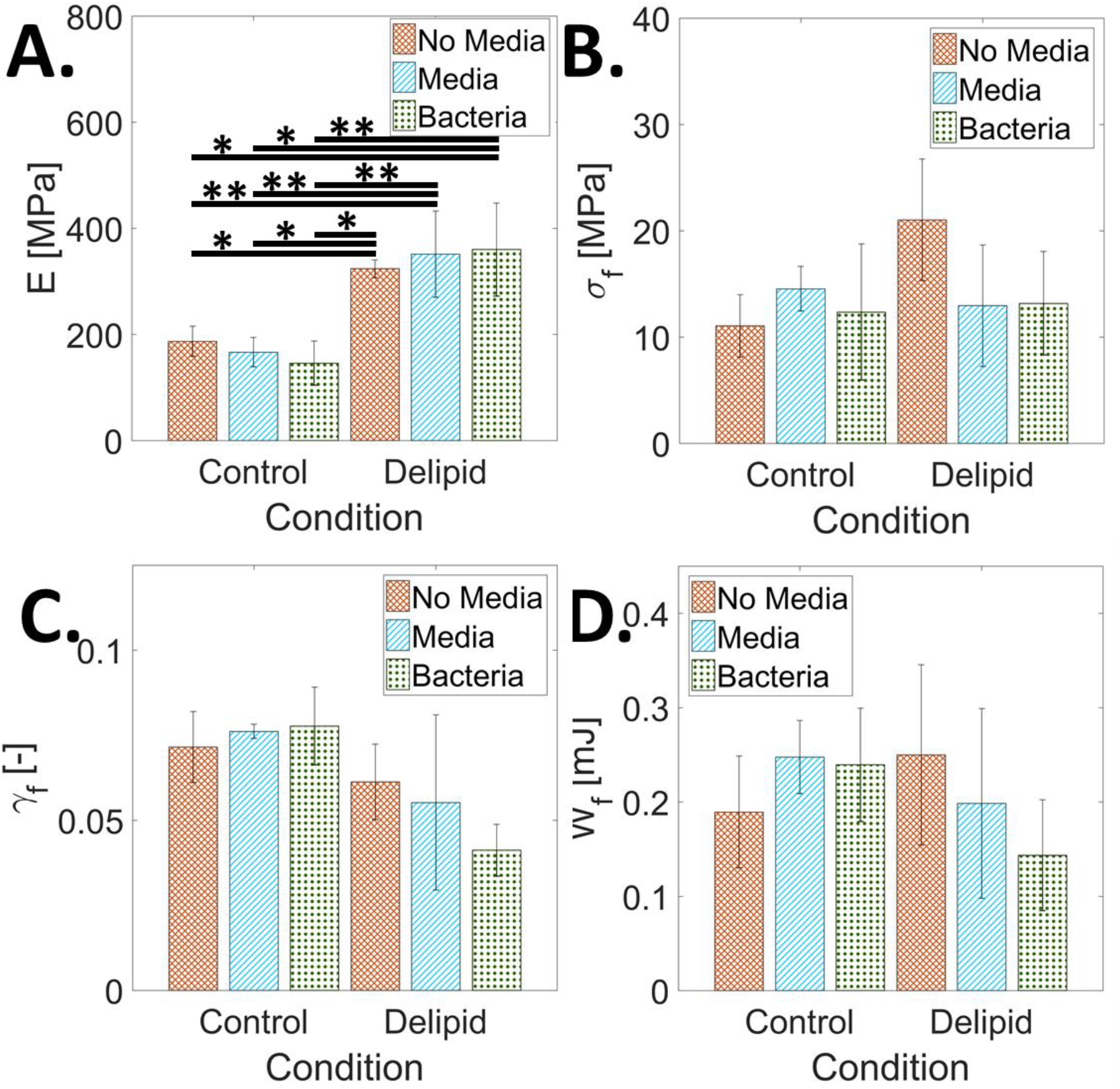
The effect of *S. aureus* tissue permeation, lipid loss, and media immersion on the mechanical properties of SC equilibrated to 25% RH. Average (A) elastic modulus, *E*, (B) fracture stress, *σ_f_*, (C) fracture strain, *γ_f_*, and (D) work of fracture, *W_f_*, for control and delipidated SC samples. Mechanical properties of SC not immersed in media are reported with orange crosshatch bars. The effect of media immersion is reported as blue striped bars. Combined media immersion and bacterial exposure effects are reported as green dotted bars. Bars denote average values of 3 ≤ *n* ≤ 8 individual sample measurements for each treatment condition. Error bars denote standard deviations.

Fig. 4 shows complementary mechanical results for SC samples equilibrated for 24 hr to 100% RH prior to mechanical testing. This conditioning enables SC tissue to undergo plastic deformation before rupture. In contrast to the 25% RH results in Fig. 3A, Fig. 4A shows that the elastic modulus does not change for delipidation alone, nor with media immersion of control SC. However, the elastic modulus does significantly increase with media immersion of the delipidated SC, and with the addition of bacteria for both lipid conditions independently. Fig. 4C similarly shows that lipid depletion alone has no significant effect on the fracture strain; however, the addition of bacteria for both lipid conditions independently causes a significant decrease in fracture strain. Supplemental Fig. S1 shows that the addition of bacteria for both lipid conditions decreases the ability of the SC to plastically deform. However, no significant effects are observed with the fracture strain with media immersion for either lipid conditions. Fig. 4B shows that fracture stress significantly increases only after lipid depletion and does not change with media immersion or the addition of bacteria for either lipid condition. Fig. 4D highlights that work of fracture significantly decreases only after the combined effects of lipid depletion, media immersion and the addition of bacteria.

**Figure 4.**
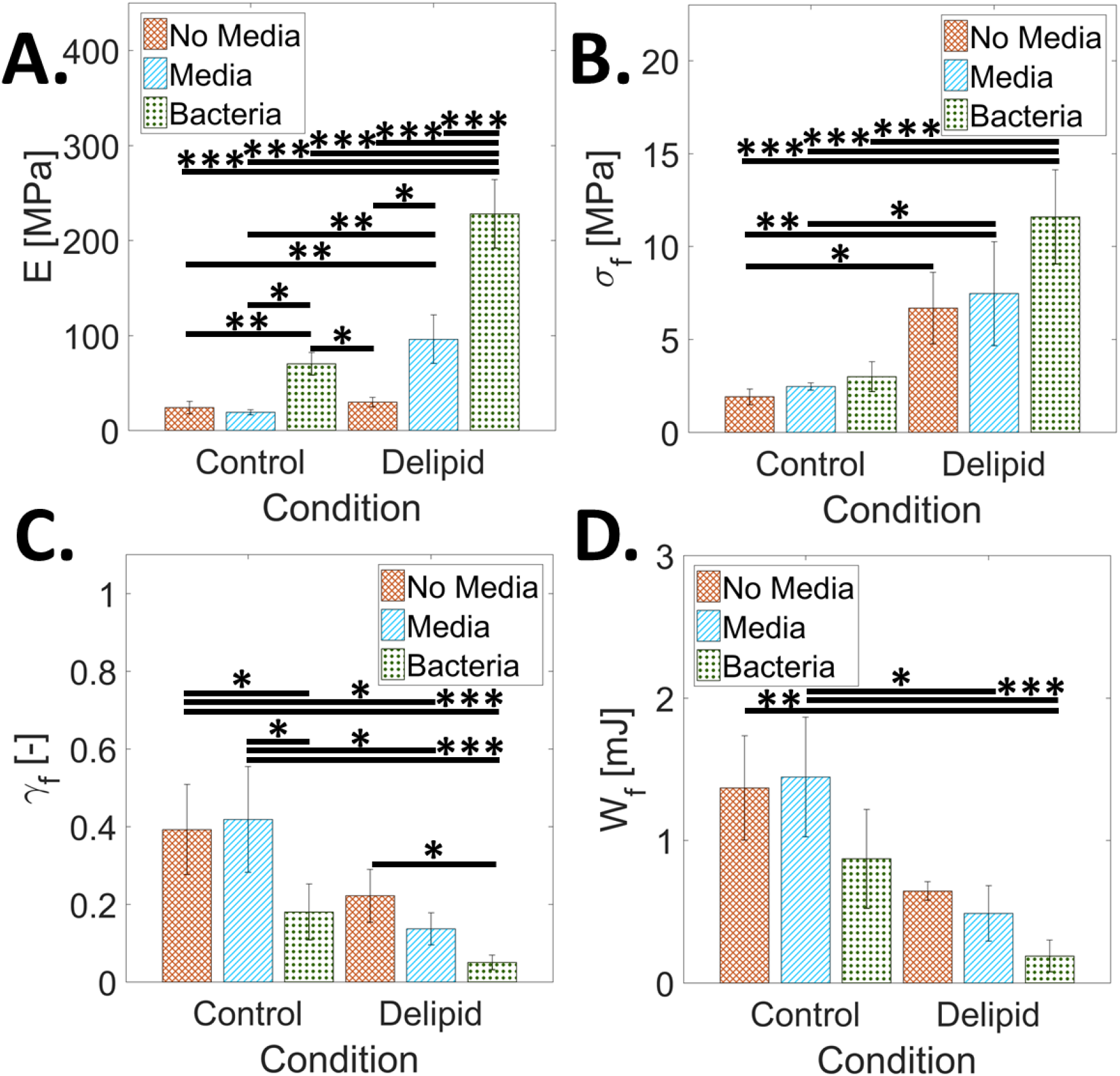
The effect of *S. aureus* tissue permeation, lipid loss, and media immersion on the mechanical properties of SC at 100% RH. Average (A) elastic modulus, *E*, (B) fracture stress, *σ_f_*, (C) fracture strain, *γ_f_*, and (D) work of fracture, *W_f_*, for control and delipidated SC samples. Mechanical properties of SC not immersed in media are reported with orange crosshatch bars. The effect of media immersion is reported as blue striped bars. Combined media immersion and bacterial exposure effects are reported as green dotted bars. Bars denote average values of 3 ≤ *n* ≤ 8 individual sample measurements for each treatment condition. Error bars denote standard deviations.

### Lipid content governs crack pathways during SC rupture

In order to better understand how delipidation and bacterial permeation induce mechanical degradation of SC, along with what constituent components of the structurally heterogeneous tissue are affected at the microscale, tissue fractography studies are performed. Here, the impact of lipid depletion, media immersion, and bacterial permeation on changes to the failure pathways that form during SC rupture are quantified. Widefield histological imaging and scanning electron microcopy (SEM) are used to identify intact corneocyte cell edges along the failure pathway, as shown in Fig. 5. Similar failure pathways are observed both at low and high humidity conditions, as well as with and without immersion for each independent lipid condition. Using histological images in Fig. 5A–D, the proportion of the failure pathway coincident with corneocyte cell edges (intercellular failure) or ruptured cells (intracellular failure) is summed and normalized by the total failure path length. Averaged values for each condition shown in Fig. 6 reveal that intercellular failure is statistically predominant for control SC conditions, while intracellular failure is significantly greater for delipidated conditions, regardless of bacterial or media exposure.

**Figure 5.**
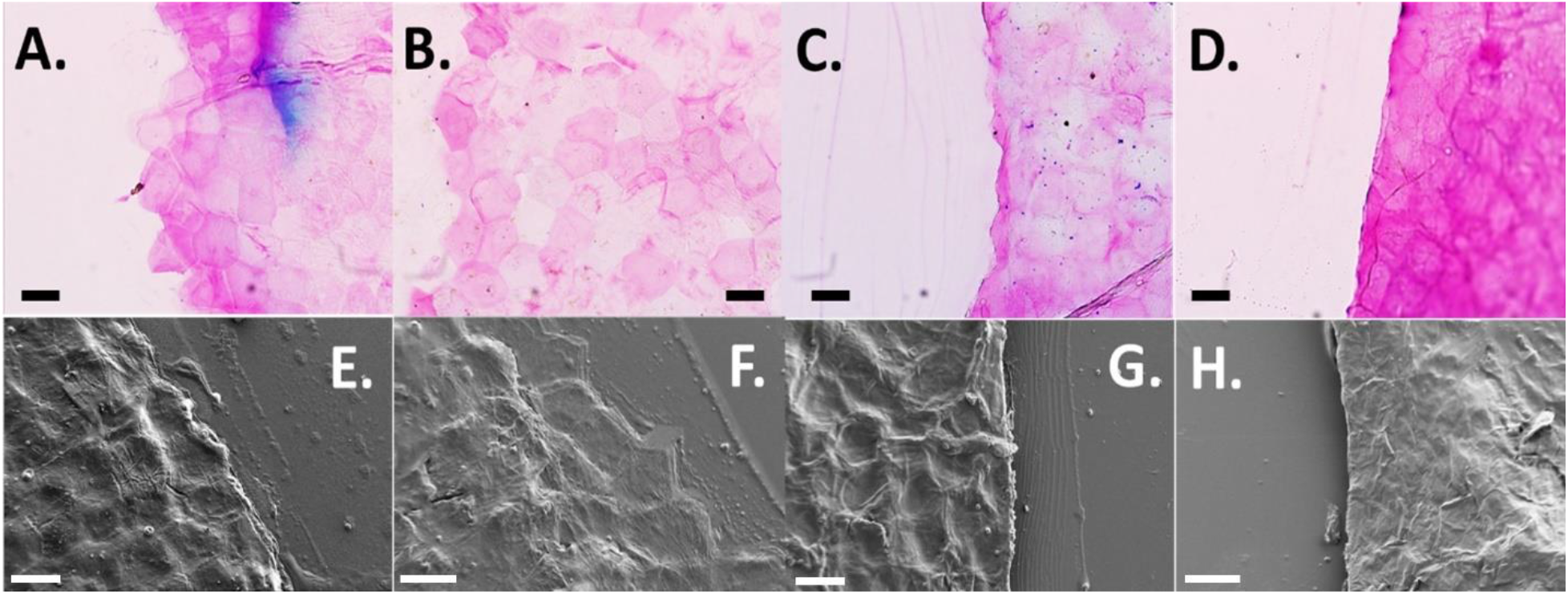
Effect of lipids and bacteria on SC failure pathway. (A–D) Brightfield images of SC crack pathways using TBBF stain to highlight polygonal shaped corneocytes. (E–H) SEM images of SC crack pathways. *Scale bars* – 30 μm. Conditions tested include control SC exposed to media only (A & E), control SC exposed to media and bacteria (B & F), delipidated SC exposed to media only (C & G), and delipidated SC exposed to media and bacteria (D & H).

**Figure 6.**
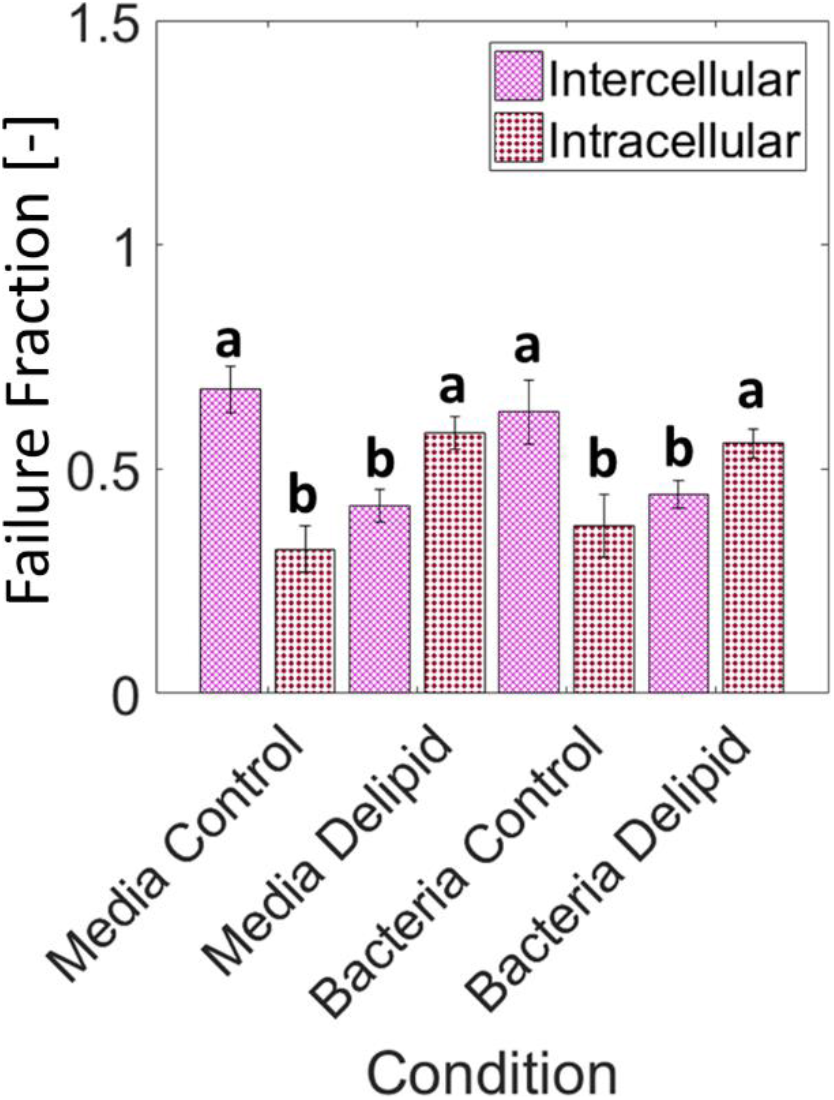
Intercellular and intracellular failure fraction during SC rupture. Average percentage of total inter- (*purple crosshatch*) and intra- (*red dots*) cellular failure, normalized by total SC crack length for different lipid and exposure conditions. Bars denote average values from *n* = 114 ± 5 individual corneocytes from *n* = 3 SC samples for each lipid and exposure condition. Error bars denote standard deviations. “a” and “b” in the figure independently denote bars that are not statistically different from one another, but are statistically different to the contrasting letter.

## DISCUSSION

Previous studies have shown that *S. aureus* can permeate into and across partially delipidated SC, with bacteria located within the tissue by day 6 of flow cell incubation at room temperature (23^°^C) and full permeation by day 9(11). Using a more physiologically relevant drip chamber arrangement that more closely mimics skin temperature and diurnal hydration/dehydration cycles, we find that permeation into the tissue occurs more rapidly. Fig. 1D shows bacterial penetration into SC by day 4. We speculate that the accelerated rate of penetration is related to the increased temperature (23 to 33^°^C), which more accurately represents average skin surface temperature (18, 19). The lack of inversion used in the flow cell setup may also increase permeation rates, due to increased initial bacterial sedimentation on the skin. Further, the bacterial presence within corneocytes only under delipidated SC conditions confirms bacterial permeation via intra and intercellular pathways, further supporting a previously published diffusion model (11) that predicts bacteria are more likely to take a direct transcellular permeation route rather than a purely intercellular pathway. Previous literature has reported the internalization of *S. aureus* into other non-professional phagocytes including osteoblasts (20–22), keratinocytes (16, 23, 24), and endothelial cells (25–27). The results of Fig. 2 indicate that bacterial internalization does occur with corneocytes, but only when the tissue is partially delipidated.

We additionally investigate the effects that bacterial colonization, penetration, and bacterial internalization have on the mechanical properties of the SC under control and partially lipid depleted conditions. At high RH, delipidation alone only increases the fracture stress. However, for both lipid conditions we observe an increase in tissue elastic modulus and a decrease in the ability to plastically deform with bacterial exposure (Fig. 4A and C respectively). In-turn, this degrades the mechanical integrity of the tissue (Fig. 4 D). A similar trend occurs with a reduction in SC water content (28–32). Previous studies report changes in the water holding capacity of SC with extracellular lipid (33) and NMF depletion (34). So while changes in the water holding capacity would explain changes in the elastic modulus of tissue due to delipidation (Fig. 3A), additional bacterial permeation may consequently also be affecting the presence of extracellular lipids (35–40) and NMFs (41–44), that will impact the water retention ability of the tissue, potentially caused by disruption of the cornified envelope during bacterial invasion into the corneocyte cells. Comparing the effect of bacteria permeation, immersion and delipidation on the elastic modulus of the SC at low RH conditions in Fig. 3A, we observe that only lipid depletion (45–49) causes a significant change in the elastic modulus of the tissue. Neither media immersion, associated with a loss of NMFs (40, 44, 50–52), nor bacterial presence impact tissue stiffness for either lipid condition.

In addition to investigating the mechanical properties of SC tissue, the impact of lipid depletion, media immersion, and bacterial growth on changes to the microstructural mechanical integrity are also quantified. When observing fracture path in control and delipidated SC tissue (Fig. 5), cracking occurs intercellularly for the control conditions, yet intracellularly for the delipidated conditions, irrespective of media immersion or bacteria growth (Fig. 6). We anticipate this lipid depletion induced change occurs due to the formation of strong interactions between adjacent lipid envelopes in intercellular regions of SC (53), resulting in an energetically less favorable failure between corneocytes and a preference for rupture of the keratin intermediate filament network within corneocytes, observable in Fig. 4B.

*S. aureus* does not produce any ceramidase(54), nor has this bacterial species previously been shown to alter stratum corneum lipids directly. Nonetheless, the impact of bacterial growth and permeation on skin shows changes in the mechanical properties of SC consistent with lipid depletion; a stiffening and weakening of the tissue. This seems therefore to suggest that the bacteria may potentially be altering other components of the tissue. The most likely alternative is that bacteria are affecting NMFs, either directly through enzyme degradation or indirectly by NMF leaching out of the cornified envelope during cellular internalization. Certainly the most prominent NMFs: Urocanic acid (UCA) (55), Ser (56), Pyrrolidine carboxylic acid (PCA) (57), Glycerol (58), and Urea (59) have all been shown to be affected directly by *S. aureus.* To ascertain the cause of bacterial medicated tissue degradation, NMF levels in SC tissue inoculated with bacteria should be compared with unexposed controls, for both normal and lipid depleted conditions, possibly using HPLC (60). It should be noted that while keratin filaments have been hypothesized to plasticize with the addition of water at high RH, contributing to SC viscoelastic properties (28, 61, 62), there have been no studies, to our knowledge, looking into the isolated mechanical effect of keratin modification in SC or the effect *S. aureus* has on keratin in skin corneocytes. These assessments should also be investigated in future studies.

## CONCLUSION

In this article we show that *Staphylococcus aureus* is internalized by corneocytes in lipid depleted human stratum corneum during permeation, indicating a viable transcellular route through the tissue. We also show the effects that changes in lipid composition, immersion, and bacterial growth into and across stratum corneum have on the mechanical deterioration of the tissue. Bacterial growth acts to weaken the tissue and make it more brittle, increasing the risk of tissue rupture. Skin suffering from atopic dermatitis is associated with depleted levels of lipids(63) and increased populations of *S. aureus* bacteria in the microbiome(64, 65). Lipid depletion and bacterial mediated mechanical degradation of the tissue could further explain the formation of exudative lesions associated with atopic dermatitis. These regions of ruptured tissue will lead to further entry of bacteria and other allergens into the viable epidermis, contributing to a chronic state of the disease. In addition, this work shows the necessity of lipids in not only preventing bacterial entry into the skin, but also governing stratum corneum mechanical barrier integrity and fracture behavior. This is especially important for people that might be prone to lipid depletion through abnormalities in lipid-processing proteins (66), occupational hazards such as repeated hand washing required for sterile environments (67, 68), or contact with metal working fluids, solvents and caustic chemicals (69). Lastly the connection between bacteria and immersion (natural moisturizing factors) could contribute to a lack of natural moisturizing factors observed in the stratum corneum of atopic dermatitis patients (70, 71).

## MATERIALS AND METHODS

### Bacterial strains

All bacterial studies used *Staphylococcus aureus* ATCC 6538 (Rosenbach, American Type Culture Collection (ATCC), Manassas, VA) isolated from human lesions. This strain was modified with *pALC2084* and edited to constitutively express green fluorescence protein (GFP). Overnight cultures were grown in brain heart infusion medium (BHI, Becton, Dickinson, Sparks, MD) supplemented with 10 mg/L chloramphenicol (Mediatech, Corning Life Sciences, Corning, NY)) for plasmid maintenance, and 250 ng/mL tetracycline (Amresco, Solon, OH) for induction of GFP, in Erlenmeyer flasks at 37 °C with agitation (220 rpm).

### Stratum Corneum isolation

A full thickness (76 yrs.) female cadaveric abdominal skin sample was obtained from ConnectLife (Syracuse, NY). In accordance to the Department of Health and Human Services regulations, 45 CFR 46.101:b:4, an exempt approval (3002-13) was attained to perform research using de-identified tissue samples. SC was isolated using standard heat bath and trypsin techniques (72). After isolation, SC sheets were placed on plastic mesh, rinsed in deionized water (DIW), and dried at room temperature and RH (23 ± 2 °C, 29 ± 3% RH).

### Stratum Corneum lipid depletion

SC sheets were divided equally into two groups: 1. a control group immersed in DIW for 60 min – this treatment does not deplete lipids or irreversibly alter the intercellular lipid structure (73), and 2. a treatment group, immersed in a 2:1 mixture of chloroform and methanol (Sigma-Aldrich, St. Louis, MO) for 60 min, partially depleting intercellular ceramides, cholesterol, and free fatty acids found in SC from the tissue (49). From estimates of human SC lipid concentrations and composition (46, 74, 75), treatments using similar solvent extraction protocols on human and porcine SC (45–48) reduce lipids by 54 ± 30%. SC samples were then punched out from both control and lipid-depleted (Delipid) SC tissue sheets with either a circular 6 mm diameter punch (Harris Uni-Core, Redding, CA) for the internalization assay (*n* = 30 control; *n* = 30 delipid) or a rectangular 9.53 × 19.05 mm punch (SYNEO, Angleton, TX) for mechanical testing (*n* = 26 control; *n* = 26 delipid).

### Substrate preparation for mechanical testing

A 20% w/v ethylene-vinyl acetate (EVA) in toluene was prepared and spin coated (WS-400B-6NNP/LITE, Laurell Technologies Corporation, North Wales, PA) on to a glass coverslip (25 × 75 mm, Electron Microscopy Sciences, Hatfield, PA) at 50 rpm for 20 sec. After evaporation of toluene for 12 hr, this produced a uniform EVA film with a thickness of 91±17 μm. EVA substrates were then placed on a hotplate (10027-028,VWR, Radnor, PA) at 60 °C to allow the EVA to soften for SC sample attachment. This temperature has shown not to create irreversible changes to the structural properties of SC(76). Control and delipidated SC samples were then alternately embedded along the long axis of the cover slip (*n* = 6 samples arranged in a 3 × 2 grid on each substrate) leaving only their outermost face exposed. This embedding process occludes the sides and underside of the SC sample, preventing bacterial growth in these regions. For each substrate tested, the order of the conditioned SC samples deposited was randomized. Substrates were then degassed in a vacuum desiccator (5310-0250, Nalgene®, ThermoFisher Scientific, Waltham, MA) with attached vacuum pump (ME4 NT Vacuubrand, BrandTech, Essex, CT) for 4 hr. This process eliminated microbubbles between the SC and EVA.

### Drip chamber setup and inoculation

Embedded SC substrates were sterilized under germicidal ultraviolet light (254 nm) for 15 min, then mounted in the custom-built drip chamber (supplemental Fig. S2). This UV treatment does not alter the mechanical properties of the tissue(77). The drip chamber is an aluminum hollow block (10×112×34 mm) that has eight samarium-cobalt magnets (M14X116DISmCo, Apex Magnets, Petersburg, WV) at each corner of the underside and top. Substrates with embedded SC samples were sandwiched between the bottom steel plate and the block’s magnets, holding the substrate in place magnetically. Vacuum grease (Dow Corning, Midland, Michigan) was also applied around the edges of the substrate to ensure the drip chamber was fully sealed. The upper steel plate holds another glass cover slip in place magnetically. This upper plate and cover slip could be slid backwards to allow for the addition of medium over the SC surface. The shorter sides of the block have threaded holes for a brass inlet (5454K62, McMaster-Carr, Chicago, IL) and outlet. The brass inlet was connected to a gas permeable filter (200 nm pore size) via silicone tubing (96400-14, Cole-Parmer, Vernon Hills, IL) to maintain atmospheric pressure. The brass outlet was connected to a cross connector (5463K93, McMaster-Carr, Chicago, IL) via silicone tubing. The cross connector separates the injection ports for the fatty acid dye (D3835, ThermoFisher Scientific, Waltham, MA) and bacterial inoculum, and an outlet port for waste. Each port on the cross connecter was connected to a Luer lock coupling (51525K322, McMaster-Carr, Chicago, IL) via silicone tubing. The drip chamber was first injected with 5 mL fatty acid dye at a 10 μM concentration in 10% BHI medium (Franklin Lakes, NJ). The lipid stain was allowed to bind to all SC samples for 30 min. To remove unbound dye from the chamber, 15 mL phosphate buffered saline (PBS) was injected into the chamber, then decanted out of the outlet port. The fluorescent fatty acid stain was found not to alter the growth behavior of the bacteria. Following staining, the chamber was injected with 5 mL of a stationary phase culture of *S. aureus* (10^8^ CFU/mL). The bacteria were allowed to attach to the SC surface for 2 hr. Unattached bacteria were washed away with 15 mL of PBS, injected into the chamber and then decanted. The top of the drip chamber was then slid backwards and 2 drops (50 μl/drop) of 20% BHI medium, supplemented with 250 ng/mL tetracycline for induction of GFP, was added to each SC sample to maintain hydration and bacterial growth. The top was then slide back into place, and the chamber was placed on a hotplate (10027-028,VWR, Radnor, PA) to simulate average skin surface temperature at 33 °C (18, 19). Media was added to the chamber each day for 5 days total until sterilization and removal of the SC from the substrate for mechanical testing. The setup was kept in a biosafety level 2 hood (Class II Type A2, Labconco, Kansas City, MO), except for imaging.

### Microscopic imaging of bacterial colonization of SC

*S. aureus* growth was monitored every 24 hr using confocal laser scanning microscopy (Leica SP5, Wetzlar, Germany) over a 5-day period. Images with a spatial resolution of 0.38 μm/pixel (1024 × 1024 pixels) were acquired using a 40x objective lens with a numerical aperture of 1.25. SC samples were illuminated sequentially with transmitted light, then at 455 and 543 nm. Images for the latter two excitation wavelengths were captured respectively across a bandwidth of 500-540 and 560-590 nm. Transmitted light images were used to distinguish topographical regions of the SC to enable consistent imaging of the same position every day, to within ~5 μm spatial accuracy. The SC position imaged was carefully chosen as to not include voids in the SC tissue arising from apocrine pores(11). The 455 nm illumination was used to excite GFP-tagged *S. aureus.* The 543 nm illumination was used to excite the fluorescent BODIPY stained SC. At each recorded time point, z-stack images were taken across the full depth of the SC sample and substrate at height increments of 0.15 μm.

### Internalization assay

Individual wells in 24-well plates (*n* = 4 plates total) were coated with 200 μl of 20% EVA polymer dissolved in toluene. The solvent was then allowed to evaporate for 12 hr. Control (*n* = 8) and delipid (*n* = 8) SC samples were then independently embedded in each well leaving only their outermost face exposed. Wells containing EVA alone (*n* = 8) were used as tissue free controls. Each 24-well plate was then degassed in a vacuum desiccator for 4 hr to eliminate microbubbles between the SC and EVA. The well plates were then sterilized under ultraviolet light for 15 min. Following this, each well was filled with 300 μl stationary phase culture of *S. aureus* (10^8^ CFU/mL). The bacteria were allowed to attach to the SC surface or EVA polymer alone for 2 hr. Unattached bacteria were washed away with three rinses of PBS. Each day thereafter, for 5 days, 2 drops (50 μl/drop) of 20% BHI medium, supplemented with 250 ng/mL tetracycline, were added to each well to maintain hydration and bacterial growth. Plates were then placed on a hotplate at 33 °C. On the 5^th^ day, methods described in Kintarak *et al* were used (16). Half of the wells in each condition (Control; Delipidated; EVA alone) were exposed to a 1 mL solution of 100 μg/mL of gentamicin, killing extracellular bacteria. The plate was then incubated for 2 hr at 37 °C and rinsed twice with PBS. Each well was then filled with 1 mL 0.5% Triton X-100 (Sigma-Aldrich, St. Louis, MO). Wells containing SC were macerated with a pipette tip. To determine internalized bacterial viable counts, serial dilutions of each well were plated on 2:1 plate count agar (247930, BD, Franklin Lakes, NJ) to standard agar (214530, BD, Franklin Lakes, NJ).

### SC sterilization, removal, and mechanical testing

After 5 days of bacterial colonization and subsequent permeation, substrates were sterilized with a 2.5 mL/L aqueous bleach solution (Clorox, Oakland, CA) in DIW for 30 mins (78–80). After sterilization, substrates were placed in a water bath at 60 °C for 5 mins to soften the EVA and dislodge the SC rectangles for subsequent mechanical testing. Supplemental Fig. S3 shows that this concentration of bleach solution did not cause significant changes in SC mechanical properties. SC samples were then equilibrated for 24 hr to either 25% or 100% RH before mechanical testing (3 ≤ *n* ≤ 8 independent samples for each lipid condition and humidity). Equilibration to low or high RH conditions was achieved by placing specimens respectively in an airtight container filled with desiccant (Drierite 10-2 mesh, W.A. Hammond Drierite Company, Xenia, OH) or a hydration cabinet (F42072-1000, Secador, Wayne, NJ) with a base filled with DIW. In both cases, RH conditions were monitored throughout the equilibration period using a hygrometer with probe (445815, Extech Instruments, Nashua, NH)). After equilibration, the mechanical properties of samples were evaluated using a uniaxial tensometer (UStretch, CellScale, Waterloo, ON, Canada) equipped with a 4.4 N load cell. The ends of each SC sample were taped (General-Purpose Laboratory Labeling Tape, VWR, Radnor, PA) to prevent slippage of the sample in the tensometer grips, leaving an exposed area of 9.53 × 10 mm. Individual SC samples were mounted into opposing tensometer grips, initially separated by 10 mm. Samples were strained until rupture at a constant strain rate of 0.012 s^−1^; similar to rates used in previous mechanical studies of skin (81). Tensile forces and grip separation were recorded at a frequency of 5 Hz. After mechanical testing, the average thickness of the ruptured SC sample was quantified with optical microscopy using an Eclipse Ti-U inverted microscope (Nikon, Melville, NY) with 40X oil objective lens (Nikon Plan UW, Nikon, Melville, NY). Images were recorded using a digital CCD camera (Andor Clara, Belfast, Northern Ireland). Optical thickness measurements were taken a distance from the crack interface to prevent measuring reduced thicknesses arising from plastic deformation. Combinations of sample dimensions, and recorded force-displacement data were then used to derive engineering stress-strain curves, from which the average elastic modulus, fracture stress, fracture strain and work of fracture were extracted. SC subjected to the same protocol without the addition of bacteria was used as media controls for each lipid condition.

### SC failure pathway imaging and failure method calculation

After mechanical testing, fractured SC sample pieces were subjected to either SEM (*n* = 3 sample halves per condition), or histological staining and widefield transmission microscopy (*n* = 3 sample halves per condition) to determine the proportion of inter- and intra-cellular crack pathways. For SEM imaging, fractured halves were laminated onto square glass coverslips (22 X22X-2, Thermofisher Scientific, Waltham, MA) and adhered to SEM pin mounts (16144, TED PELLA, Redding, CA) using carbon tape (16084-3, TED PELLA, Redding, CA). To increase conduction, copper tape (102091-345, VWR, Radnor, PA) was used to create a path from the bottom of the chuck to the fractured SC sample. The SEM substrate was then sputtered using a carbon coater (208C High Vacuum Turbo Carbon Coater, Cressington). SC failure pathways were then imaged on an SEM (ZEISS FEG-SEM Supra 55 VP) with a voltage of 0.7–1.5 kV. For histological staining, fractured halves were laminated onto glass slides (16004-422, VWR, Radnor, PA) and exposed to toluidine blue and basic fuchsin (TBBF) stain in 30% ethanol (Delasco, Council Bluffs, IA) for 1 min. This process dyes the intercellular regions of the SC, enabling visualization of individual corneocytes. Slides were then rinsed with DIW for 1 min and allowed to dry for 2 hr. A drop of mounting media (8310-16,Thermofisher Scientific, Waltham, MA) was then added to each slide before the coverslip was mounted (48393-059, VWR, Radnor, PA). Prepared slides were then imaged along the failure pathway using a light microscope (BX43, Olympus, Tokyo, Japan) and 50x objective.

Images of the SC failure pathway from the TBBF stained tissue samples were used to quantify the total fracture crack length and the proportions of both intercellular (Fig. 7A) and intracellular failure (Fig. 7B). Intercellular failure resulted with intact corneocytes with well-defined polygonal edges. Intracellular failure resulted with broken corneocytes without well-defined polygonal morphologies. Images were taken interspersed across the entire width of the SC sample (9.53 mm), representing 13-21% of the total crack length (*n* = 3 samples per condition, *n* = 114 ± 5 corneocytes total per condition). Corneocytes with edges that were not all well distinguished were not used, as shown in Fig. 7C. The total lengths of intercellular and intracellular failure normalized by the total crack length measured provide the proportion of intercellular to intracellular failure for each condition.

**Figure 7.**
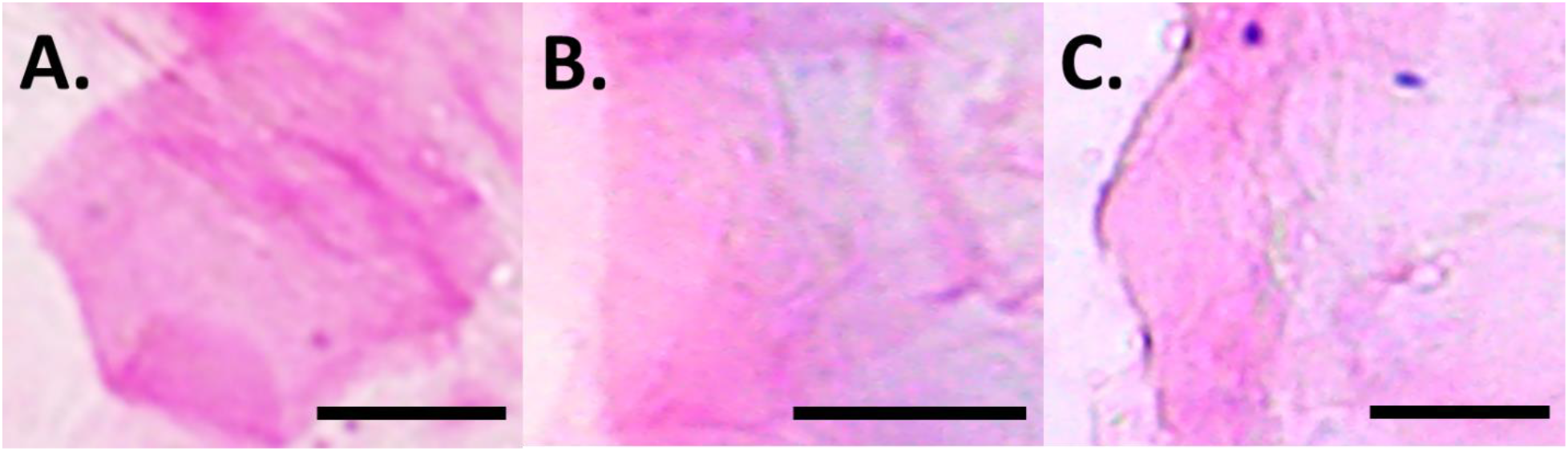
Corneocyte integrity along SC failure pathway. Examples of TBBF stained (A) intact, (B) broken, and (C) indistinguishable corneocytes imaged at 50x along a fractured SC pathway. *Scale bar* – 15 μm.

### Statistical analysis

All statistical analyses were performed using R (version 3.4.2). A 1-way ANOVA was used to test for statistical significance in: Fig. 2, where the addition or lack of gentamicin was compared independently across all sample types (Control, Delipid, and EVA alone), Fig. 3 and Fig. 4, where comparisons were made across all lipid and exposure conditions, and Fig. 6, where comparisons are made across all failure preferences, and lipid and exposure conditions. Levene’s and Shapiro-Wilk’s tests were respectively used to determine equality of variances and normality. Results in Fig. 2A and B were found to exhibit non-normal distributions and unequal variance. Here a Kruskal-Wallace analysis was performed. Results in Fig. 3A and Fig. 4B and C were found to exhibit normal distributions and unequal variances. Here a 1-way ANOVA with Welch correction was performed. Results in Fig. 3B–D, Fig. 4A and D, and Fig. 6 were found to exhibit normal distributions and equal variances. Here a standard 1-way ANOVA was performed. Post-hoc analyses were performed if statistical significance levels below 5% were established. In the figures, * denotes *p* ≤ 0.05, ** denotes *p* ≤ 0.01, *** denotes p ≤ 0.001, and matching letters denote *p* ≥ 0.05.

## Abbreviations

AD: Atopic Dermatitis
SC: Stratum corneum
EVA: Ethylene-vinyl acetate
RH: Relative humidity
SEM: Scanning electron microscopy
ClfB: Clumping factor B
FnBPB: Fibronectin binding protein B
NMFs: Natural moisturizing factors
UCA: Urocanic acid
PCA: Pyrrolidine carboxylic acid
GFP: Green fluorescent protein
BHI: Brain heart infusion
DIW: Deionized water
PBS: Phosphate buffered saline
TBBF: Toluidine blue and basic fuchsin

## ACKNOWLEDGEMENTS

This material is based upon work supported by the National Science Foundation under Grant No. 1653071 and Grant No. 1757846. We would also like to thank Elana Chazen, Amanda Kenny, and Caitlin M Begley for originally designing the drip chamber device. We would also like to thank F. D. C. Willard for useful conversations about this work.

